# Bioimaging Data Management Workflow for Plasma Medicine

**DOI:** 10.64898/2026.01.26.700509

**Authors:** Mohsen Ahmadi, Robert Wagner, Sander Bekeschus, Markus M. Becker

## Abstract

Bioimaging experiments in plasma medicine generate complex datasets that go beyond conventional imaging studies by combining microscopy data with heterogeneous metadata from biological experiments and highly parameterized gas plasma treatments. Plasma exposure conditions, such as device configuration, gas composition, and treatment conditions, are critical determinants of biological outcome, yet they are rarely captured in a standardized, machine-readable, and reusable manner. To address this gap, we present a research data management (RDM) workflow that operationalizes the Findable, Accessible, Interoperable, and Reusable (FAIR) principles across the bioimaging data lifecycle in plasma medicine. The workflow is implemented as a structured pipeline integrating open-source tools, including OMERO for image data management, eLabFTW as an electronic laboratory notebook, Adamant for schema-driven metadata collection, and Micro-Meta App for standardized documentation of microscopy acquisition settings that are connected via programming interfaces to enable persistent linkage of metadata to image datasets using standardized annotations. The workflow is documented in a reproducible tutorial with an open-source Python Jupyter notebook hosted on GitHub. By integrating plasma treatment metadata with imaging data, this approach improves reproducibility, cross-study comparability, and data reuse in plasma medicine research.

## Introduction

Plasma medicine is an interdisciplinary field that explores the application of cold gas plasma (a partially ionized gas containing, e.g., free electrons, ions, and reactive species ^1^) for therapeutic and biomedical purposes ^2,3^. It combines principles from plasma physics, biology, medicine, and engineering to develop novel treatments and diagnostic tools for various diseases. Microscopy-based imaging is one of the main pillars of experimental research in plasma medicine, offering insights into how cells and molecules respond to plasma treatment, which guides the study of gas plasma-induced biological effects ^4-6^. These processes are dynamic and often require quantitative time-resolved imaging to capture changes across varying conditions, exposure times, and biological models, thereby supporting a mechanistic understanding of plasma-target interactions. As image acquisition technologies advance toward high-throughput and high-content formats, establishing structured and reproducible bioimage workflows has become essential for integrating plasma medicine research into the global bioimaging community, which is actively working to standardize metadata practices ^7,8^. However, research data management (RDM) in microscopy is challenged by the large volume and heterogeneity of the image data. Experimental records, raw imaging files, and metadata are often stored in separate systems, such as laboratory notebooks for protocols and metadata, and image repositories, such as OMERO, for raw data ^9^. This fragmentation hinders reproducibility, transparency, and traceability, and ultimately limits compliance with the F(findable)A(accessible)I(interoperable)R(reusable)-data principles ^10^. By establishing semantic linkages between raw data, metadata, and analytical outputs across experimental stages, an image data management workflow lays the groundwork for interoperable RDM infrastructure in imaging-related research in plasma medicine.

Microscopy plays a central role in plasma medicine by enabling the detailed visualization of how cells respond to plasma-induced effectors, such as reactive oxygen and nitrogen species (ROS/RNS). These ROS/RNS can influence key biological processes, such as apoptosis, cell proliferation, and oxidative signaling. To investigate these effects, researchers commonly employ fluorescence microscopy ^11^, live-cell imaging in multi-well formats ^12^ (e.g., time-lapse imaging of multiple conditions within a single plate), drug screening and dose-response assays across defined compound panels ^13^, and automation-assisted fluorescence imaging ^14^ of multi-well plates, where individual wells correspond to specific treatments, genetic perturbations, or experimental conditions. Such automation provides systematic image acquisition across many wells while maintaining controlled experimental variability, but only becomes high-content imaging or screening when combined with multiparametric feature extraction and large-scale automation ^15^. These complications create a unique RDM challenge, as multiple large sets of structured and unstructured data, including images, protocols, reagents, treatment parameters, and analysis outputs, must be consistently linked, contextualized, and made accessible. These datasets may be shared via the BioImage Archive (BIA: https://www.ebi.ac.uk/bioimage-archive/) or Image Data Resource (IDR: https://idr.openmicroscopy.org/).

Despite its importance, plasma medicine lacks FAIR-aligned RDM practices tailored to imaging to overcome the traditional RDM limitations, such as unstructured metadata, manual and error-prone data handling, poor linkage between images and experimental parameters, and the use of proprietary or non-standardized formats (Fig. 1). As shown, image acquisition and experimental metadata collection occur separately, resulting in heterogeneous and often incomplete metadata sets. Image analysis generates derived data that are not systematically linked to raw images. Local storage functions as a central repository but lacks standardization, version control, and traceability. In this context, several tools have been developed to address the broader challenges of microscopy data handling. OMERO is one of the most widely adopted solutions, providing centralized data storage, access control, metadata annotation, and integration with analysis workflows, capabilities that benefit facility staff and end users ^16^. OMERO is widely adopted by imaging facilities ^16^ to standardize metadata handling, data organization, and reproducible workflows, demonstrating its applicability for institutional-scale bioimaging data management and sharing ^17^. Importantly, public image-data repositories, such as the IDR, are built on the OMERO infrastructure and APIs, providing public submission, query, and access via the OMERO/web API ^18^. The BIA supports broad life-science microscopy data deposition and uses the OMERO ecosystem and OME formats as a key part of its ingestion and metadata workflows ^19^. These cases show that with the right setup, OMERO can transform the handling of imaging data across research institutions and public repositories.

**Figure 1.**
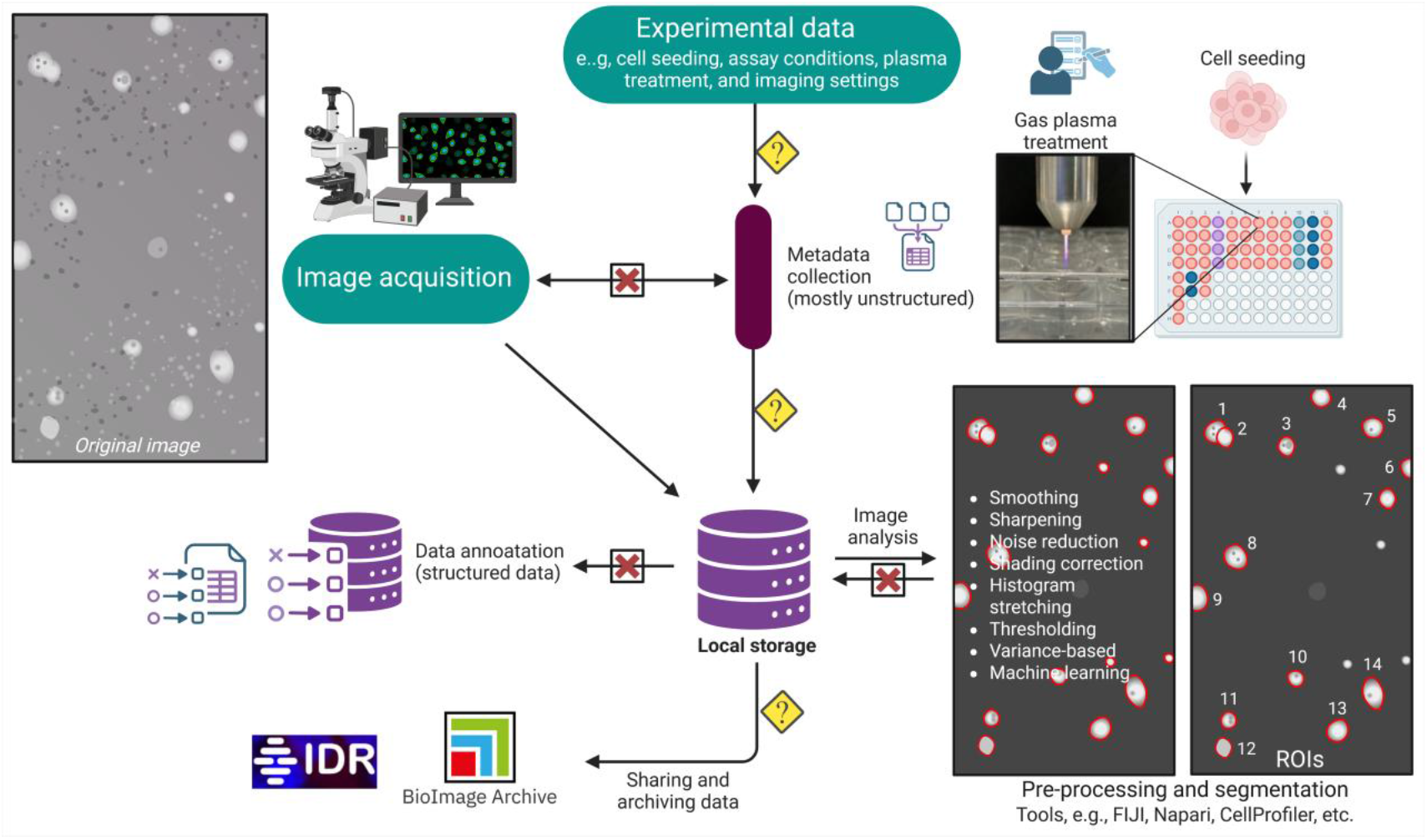
Overview of a traditional example of local bioimaging data management. Image acquisition produces primary raw image data along with the experimental parameters (cell seeding and gas plasma treatments) captured both during setup and during imaging, yet the associated metadata collection remains largely unstructured. Data are typically stored locally, annotated manually, and processed using analysis tools. The resulting datasets, often fragmented across files and folders, may create problems with sharing and archiving open repositories such as IDR and BIA. (X) Missing or manual data transfers; (?) Indicate uncertainties, such as incomplete metadata and unclear storage procedures. Created in BioRender. Bekeschus, S. (2026) https://BioRender.com/6tae47g.

Building on OMERO’s capabilities, Massei et al. ^20^ developed semi-automated workflows that combined OMERO with workflow management systems to align HCS bioimaging data management with the FAIR principles. Complementary to this, Escobar Díaz Guerrero et al. ^21^ introduced LEO, a platform designed to link OMERO with electronic laboratory notebooks (ELNs) and other metadata. LEO provides retrospective integration of the experimental context with imaging datasets (metadata from ELNs and external systems are connected to image data after acquisition), supporting richer contextualization.

In addition to these integration platforms, a range of specialized tools supports the standardization of individual (meta)data components commonly encountered in bioimaging workflows. Micro-Meta App supports the consistent documentation of microscope hardware configurations and acquisition settings ^22^. ELNs such as eLabFTW ^23^ (https://www.elabftw.net/), openBIS (https://openbis.ch/), and LabArchives (https://www.labarchives.com/) support standardized documentation of experimental procedures. The open-source software Adamant extends this coverage to the entire bioimaging workflow (from sample preparation and treatment conditions to data capture), providing structured and searchable records that complement imaging metadata ^24^. Furthermore, Jupyter notebooks are widely used for transparent and reproducible data processing, enabling human-readable and machine-actionable analytical workflows ^25-27^.

To achieve an applicable RDM framework for plasma medicine, these components must be integrated into a unified workflow that reflects the complexity of biological imaging experiments. Several initiatives have emerged to address these challenges. The NFDI4Bioimage consortium (https://nfdi4bioimage.de/home/) is developing a national infrastructure to support standardized and interoperable bioimage data across institutions ^28^. The Recommended Metadata for Biological Images provides tiered guidelines for reporting metadata in biological imaging ^29^, and minimum information guidelines now support standardized documentation for modalities such as 3D microscopy, multiplexed tissue imaging, and cell migration experiments ^15,30^. Plasma-MDS ^31^ is a plasma-specific metadata schema integrated into NFDI4Bioimage ^32^ to enable domain-relevant metadata annotation, including for imaging-based studies. Despite these advances, adoption in plasma medicine remains limited, with imaging workflows often relying on inconsistent documentation and ad hoc data handling that hinder alignment with the FAIR principles. To address this gap, we introduce a FAIR-aligned workflow integrating OMERO, eLabFTW, Adamant, and Micro-Meta App to link experimental (biological and gas plasma treatments) context, instrument settings, and image data within a unified and reproducible framework.

## Methods

### Defining the target group and workflow approach

To develop a user-friendly workflow, we first examined our target group to understand how bioimage data are generated in practice. The aim was to provide a workflow that meets the needs of the target users and encourages adoption. Our target users are scientists who culture and prepare tissues or cells, apply plasma treatments as an experimental intervention, and perform bioimaging before and/or after treatment using microscopy-based methods. They possess deep domain knowledge but often lack experience in RDM. The workflow focuses on the structured storage of image data and linking the collected metadata directly to the images via annotations. These annotations should be automated where possible, yet remain transparent and easy to understand. To keep the information manageable, metadata are organized across different levels, allowing users to access only the details relevant to their roles, as defined by the team leads and project scope.

### Metadata collection and schema management

Metadata collection in the workflow is based on standardized, schema-driven approaches to ensure compliance with the FAIR principles. Metadata handling and schema development are supported by Plasma-MDS (https://github.com/plasma-mds), which provides a coordinated collection of open resources for RDM in low-temperature plasma science. Plasma-MDS is a structured JSON schema-based standard defining the description of plasma experiments, including all relevant device and process parameters, and diagnostics. REMBI provides tiered guidelines for reporting metadata and controlled vocabularies in biological imaging ^29^. Metadata is captured and validated as JSON files that are stored and linked to experimental records in eLabFTW. Schema definitions and metadata templates are version-controlled and support the integration of schema-driven metadata acquisition (e.g., REMBI-compatible Excel templates for biological metadata collection) into routine laboratory practices.

### Technical infrastructure and software components of the proposed framework

The workflow was implemented using open-source software components. OMERO (https://github.com/ome/omero-server) serves as a central platform for the storage, organization, and access of microscopy images. It is complemented by OMERO.insight (https://github.com/ome/omero-insight) and OMERO.web (https://github.com/ome/omero-web) for the interactive data management, visualization, and annotation.

Adamant (https://github.com/plasma-mds/adamant) was used for schema-driven metadata collection and validation. Adamant enables the creation of structured, machine-readable metadata based on community standards and supports consistent metadata entries through customizable templates. eLabFTW (https://github.com/elabftw/elabftw) functions as an electronic laboratory notebook (ELN), providing human-readable experimental documentation and serving as a central repository for experimental records and associated metadata files generated within the workflow. The Micro-Meta App (https://wu-bimac.github.io/MicroMetaApp.github.io/) was used to document microscope hardware configurations and acquisition parameters in a standardized and interoperable manner.

All workflow components are publicly available through GitHub repositories, with the complete workflow, along with configuration files, metadata templates, and processing scripts, version-controlled and hosted at GitHub (see Code availability section). The execution of the workflow requires the configuration of host addresses for the OMERO and eLabFTW instances in the provided Python scripts (available in the script directory of the repository), which handle authenticated data transfer and metadata synchronization between the two systems.

### Workflow implementation in a Jupyter Notebook

The workflow was implemented using a Jupyter notebook to integrate the executable code with a step-by-step tutorial for users. The notebook was developed using Python version 3.12.5 and executed in the JupyterLab environment (https://jupyterlab.readthedocs.io/en/latest/).

### FAIR Implementation and Assessment

The FAIR principles, as formulated by Wilkinson et al. ^33^, provide a conceptual framework for improving the management, accessibility, and reusability of research assets. These principles form the basis for the workflow developed for the plasma medicine domain and were systematically applied to microscopy image data and associated metadata. Table 1 summarizes the FAIR principles and highlights their key aspects relevant to the practical implementation in plasma medicine. The FAIR compliance of the implemented workflow was assessed in a structured manner using the Common DSW Knowledge Model (version 2.6.12; https://registry.ds-wizard.org/knowledge-models/dsw:root:2.6.12), which maps questionnaire domains that map the technical and organizational aspects of data management to the individual aspects of the FAIR principles.

**Table 1.** Overview of the FAIR guiding principles with illustrative examples relevant to plasma medicine.

| Id | Principle | Implementation in plasma medicine |
| --- | --- | --- |
| **F** | **To be Findable** |  |
| F1 | (Meta)data have a globally unique and persistent identifier | Assign DOIs to datasets (e.g., image collections of plasma-treated cell lines and their corresponding treatment metadata). Individual files (e.g., CZI, TIF, CSV, TXT) remain versioned and traceable. |
| F2 | Data described with rich metadata (defined by R1) | Collect and capture plasma source parameters, treatment and environmental conditions, imaging acquisition settings, and biological context (cell culture conditions pre- and post-treatment) |
| F3 | Metadata includes dataset identifiers | Ensure metadata in OMERO or INPTDAT references the dataset DOI, e.g., dataset DOI linked to plasma device log. |
| F4 | (Meta)data are registered or indexed in a searchable resource | Upload datasets to Zenodo/INPTDAT/OMERO, to ensure plasma studies are searchable within repositories relevant to plasma medicine and the life sciences. |
| **A** | **To be Accessible** |  |
| A1 | Retrievable via a standardized communication protocol | Provide access via open protocols, e.g., HTTP(S), APIs for OMERO, and DOI services. |
| A1.1 | Protocol open, free, universally implementable | Use HTTP(S) endpoints without proprietary barriers; metadata formats such as OME-XML, OME-TIFF, CSV, JSON, and TXT must be used with open accessibility. |
| A1.2 | The protocol allows authentication when needed | Use logged-in access, e.g., institutional login and token-based, for sensitive pre-publication plasma and bioimage data. |
| A2 | Metadata remains accessible even if the data are removed | Keep gas plasma and biological metadata, e.g., treatment descriptions, plasma settings, cell culture conditions, and device configuration, persistent on the data repository even after lift or archive changes. |
| **I** | **To be Interoperable** |  |
| I1 | Use formal, accessible, shared language | Use OME-XML and OME-TIFF (for microscopy), ISA-Tab (experimental metadata), structured data exchange formats, e.g., JSON, JSON-LD, or Resource Description Framework (RDF), and standardized plasma parameter vocabularies where available. |
| I2 | Use vocabulary following FAIR principles | Controlled terms for plasma type, gas flow rate, cell line identifiers, e.g., use plasma-specific ontologies and biomedical standards (Plasma-MDS, ISA, MIHCSME, and REMBI) where available. |
| I3 | Include qualified references to other (meta)data | Link plasma exposure parameters with imaging metadata and cell-culture documentation to ensure each dataset is tied to its experimental context. |
| **R** | **To be Reusable** |  |
| R1 | Richly described with relevant attributes | Document all parameters required to reproduce the plasma treatment or biological experiment, including nozzle distance, ambient conditions, medium volume, and imaging settings. |
| R1.1 | Released with a usage license | Provide CC-BY licensing for plasma bioimage metadata and datasets. |
| R1.2 | Detailed provenance provided | Record the full experimental workflow, e.g., plasma source calibration, treatment sequence, and imaging pipeline. Document who generated the data, when and where it was produced, the software versions used (OMERO, microscopy software, ImageJ/FIJI, Python packages), any data transformations (e.g., CZI to TIFF), and key workflow steps such as OMERO import, annotation, and segmentation. |
| R1.3 | Meet community standards | Follow OME data formats for imaging, plasma metadata templates in Plasma-MDS, and plasma medicine reporting standards. |

## Results and Discussions

### Bioimage Data Management Workflow

The data management workflow for handling bioimaging data and metadata in plasma medicine research operates across three interconnected levels ^34^: metadata, data handling, and data processing.

#### I. Metadata

The experimental context, microscope setup, and image acquisition information are systematically documented using eLabFTW, Micro-Meta App, and Adamant. This system enables the standardized capture of metadata and linkage to image datasets.

#### II. Data handling

OMERO serves as a central platform for image management. It secures batch uploads, integrates metadata from eLabFTW, and ensures long-term data accessibility.

#### III. Data processing

The workflow enables batch analysis of fluorescence images from multi-well plates and supports customizable image analysis pipelines, allowing flexible adaptation to specific experimental requirements.

Fig. 2 illustrates these three levels with their software components and the information flow between the levels. The entire workflow is provided as a Jupyter notebook (see Code Availability section), which connects all tools by direct links and Python programming scripts, providing the immediate use of the workflow by scientists in the laboratory. The three levels and their respective technical components of the workflow are described in the following sections.

**Figure 2.**
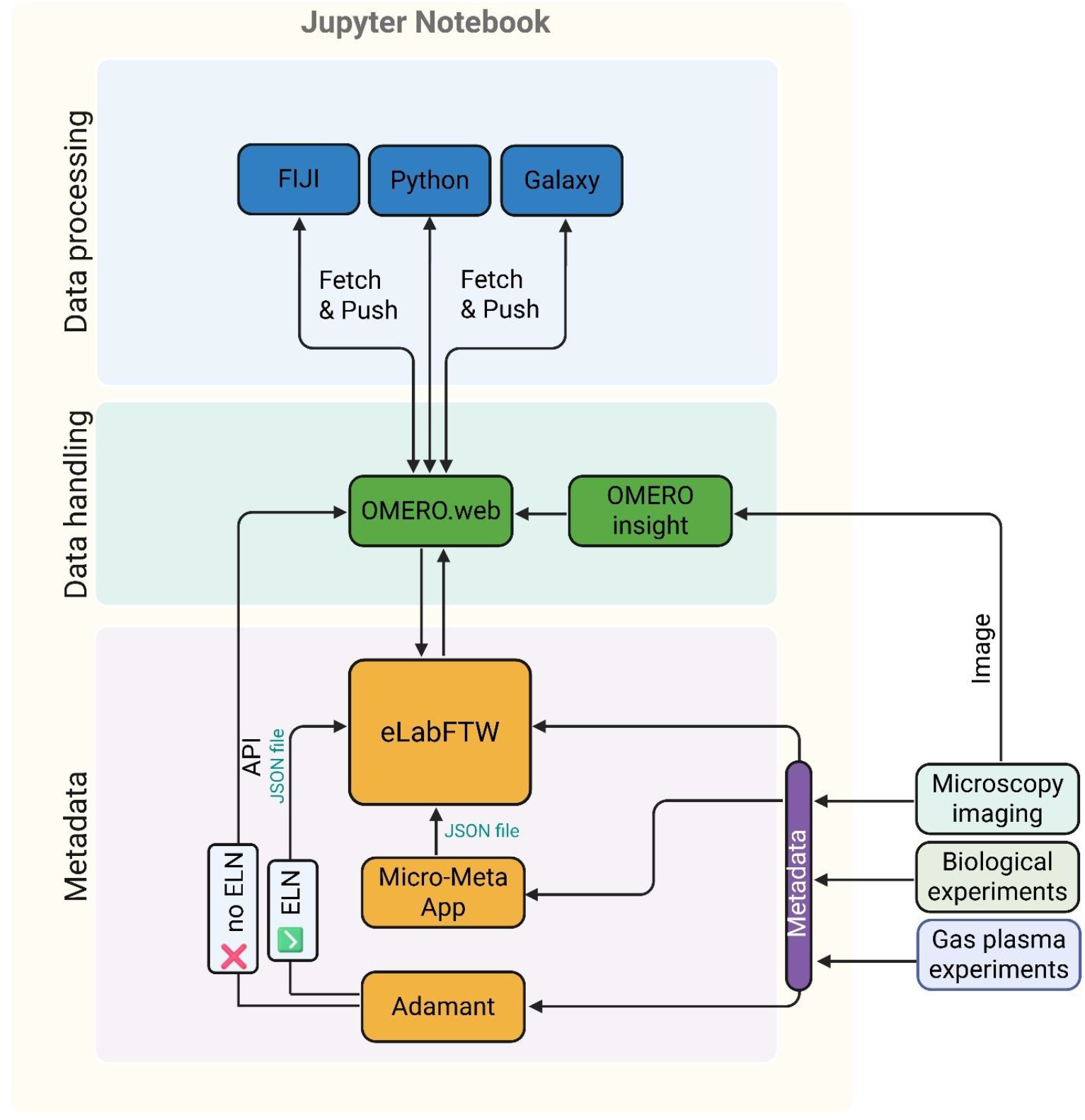
Overview of the bioimaging data management workflow for plasma medicine. The workflow is structured across three levels: metadata (Micro-Meta App, eLabFTW, and Adamant), data handling level (OMERO), and data processing level (annotation and analysis). Created in BioRender. Bekeschus, S. (2026) https://BioRender.com/1s1dfpy. A Jupyter notebook connecting all workflow components via Python scripts and user interfaces is available at https://github.com/INP-SDT/Bioimaging-Data-Management-Workflow-for-Plasma-Medicine.

### Metadata

As the first stage of the workflow, a structured metadata layer is established to capture all experimental aspects in a reusable and machine-actionable way. Metadata is systematically recorded across the entire experimental life cycle, including experimental planning and objective definition, protocol specification, sample preparation, plasma treatment, downstream biological assays, and microscopy imaging. Fig. 3 shows an overview of the metadata level of the workflow, illustrating the collection and integration of metadata from microscopy images, microscope hardware, acquisition settings, and experimental records encompassing biological assays and gas plasma treatments.

**Figure 3.**
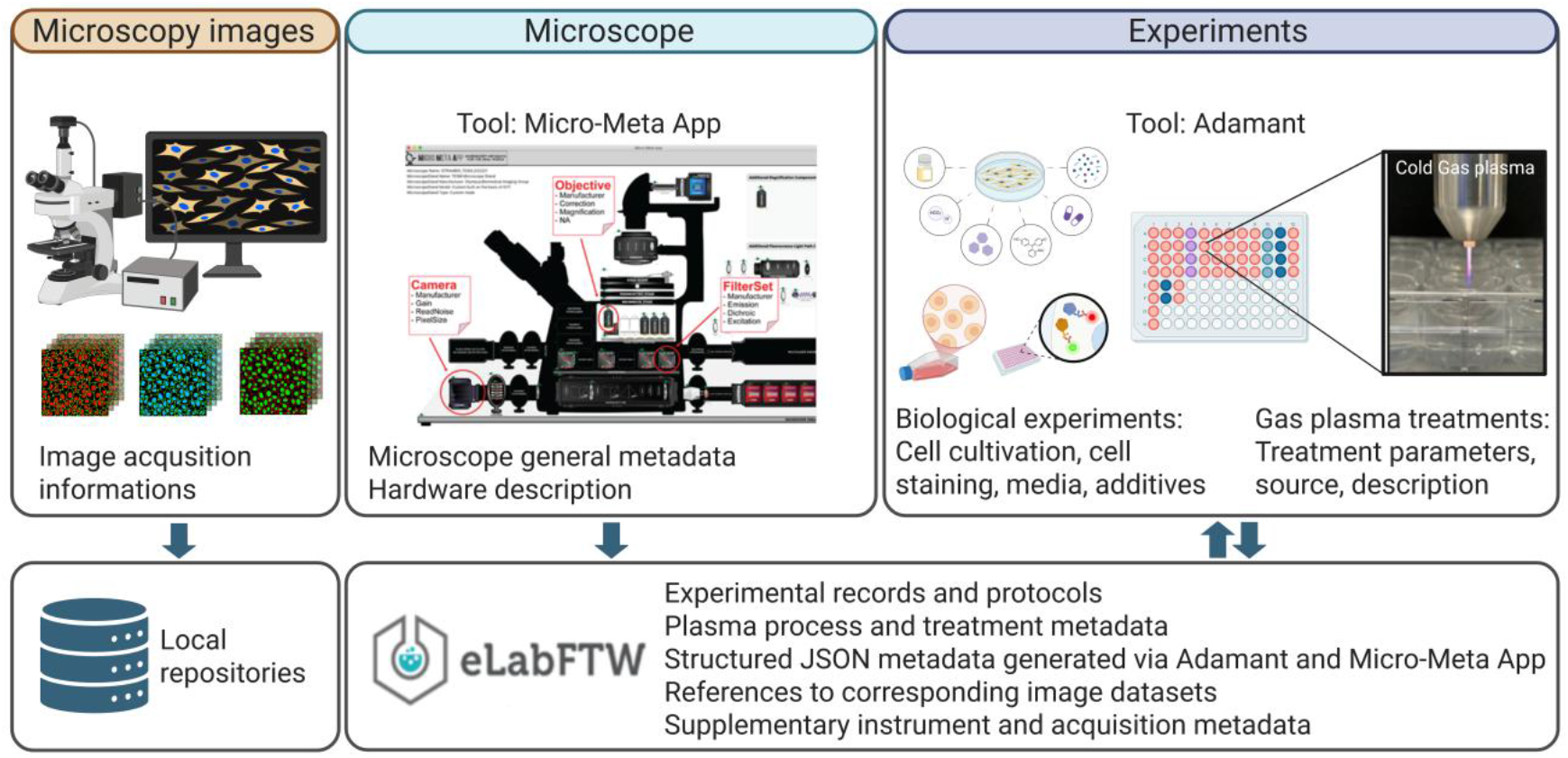
Overview of the “Metadata” level of the workflow, showing the collection of metadata from three complementary sources: (i) microscopy images, (ii) microscope hardware and acquisition settings, and (iii) experimental records encompassing biological assays and gas plasma treatment. All collected metadata are systematically stored and linked in eLabFTW across imaging and experimental workflows. Created in BioRender. Bekeschus, S. (2026) https://BioRender.com/qlvnze6.

In the workflow, the experimental context is documented as protocols and metadata details in **eLabFTW** (Fig. 2). Here, metadata are organized into three complementary categories (Fig. 3): (a) Biological experiments included detailed logging of reagents and additives (e.g., dyes and chemical treatments) with information on the source, lot number, and concentration. Biological materials (e.g., cell lines or primary cells) are annotated with provenance, passage numbers, and culture conditions. (b) Gas plasma treatment covers plasma-specific parameters such as exposure duration, gas composition, and environmental conditions during treatment. (c) Microscopy imaging documents microscope hardware specifications and image acquisition parameters, including objective lens, exposure time, detector type and sensitivity (e.g., gain and binning), laser wavelength and power (for confocal/fluorescence), and Z-stack spacing or time intervals (for 3D/time-lapse imaging). The experimental hierarchy is defined as follows: “Screen” shows the overall experiment or study, typically comprising multiple plates that together define the complete experimental campaign (e.g., plasma treatment or screening study). “Plate” corresponds to a single physical multi-well plate (e.g., a 96- or 384-well plate) within a screen, defining the layout and grouping of the experimental conditions. A “Well” represents an individual experimental unit within a plate, where a specific biological sample, treatment condition, and imaging configuration are applied and recorded.

The recorded experimental metadata in eLabFTW is used by Adamant for structured metadata generation, except for the microscope hardware specifications, which are captured separately using the Micro-Meta App. Biological experiment metadata are systematically collected using a **REMBI-compatible Excel template** and subsequently provided to Adamant as an attached file. This file is a structured metadata table used to document “Well”-level biological, imaging, and treatment information for multi-well bioimaging experiments in the context of the corresponding “Plate” and “Screen.” It captures the identification (including well position and descriptive title), cell state at the time of imaging, and detailed per-channel imaging information. For each imaging channel, the template records the channel label, signal or contrast mechanism, and targeted biological entity. In addition, extrinsic variables describing the applied treatments, including the treatment type, duration, solvent, concentration, and applied volume, together with the overall treatment regime, are documented. Plasma-specific metadata, such as plasma source name, plasma usage, cell treatment time, and treatment conditions, are captured in accordance with Plasma-MDS. Overall, this process provides metadata that links the biological state, imaging configuration, and plasma or chemical treatments for structured annotation and FAIR-compliant integration with downstream systems, including Adamant, eLabFTW, and OMERO (Fig. 2).

In the next step, structured “Screen”- and “Plate”-level metadata are collected via **Adamant** using a dedicated metadata schema (Fig. 3). The metadata schemas are aligned with established standards, including REMBI ^29^, ISA (Investigation, Study, Assay) ^35^, MIHCSME (Minimum Information for High Content Screening Microscopy Experiments) ^15^, and Plasma-MDS ^31^, thereby supporting FAIR-compliant integration with OMERO and eLabFTW software. This included unique identifiers for the “Screen” and “Plate” constructed from institutional naming conventions and linked to the corresponding eLabFTW experiment identifiers, as well as titles that matched the respective objects in the OMERO server. Descriptive metadata, such as author, description, protocol reference, and experimental dates, are recorded to provide contextual information. At the “Plate” level, biological and experimental descriptors are captured, including organism and cell type, accession numbers, and reference databases (e.g., ATCC or DSM), phenotype and genotype, culture conditions (media type, volume, concentration, and pH), seeding density, and plate-specific features such as plate format, material, and well configuration. Additional metadata includes the experiment type, imaging conditions (e.g., use of a lid, microscope name), plate preparation date, and creator information. Finally, the REMBI-compatible Excel file is uploaded and linked within Adamant, enabling the integration of “Screen”, “Plate”, and “Well” metadata into a unified, structured representation. The resulting JSON file created by using Adamant is then transferred to the ELN via the eLabFTW application programming interface (API) ^23^, where it is stored as an attachment to the corresponding experiment record (Fig. 2). Before storage of the JSON files in eLabFTW, Adamant validates the aggregated metadata against the defined JSON schema to minimize collection errors and ensure structural consistency.

Technical metadata related to imaging instrumentation is collected using the **Micro-Meta App**, a community-driven tool designed to standardize microscope descriptions and operating conditions ^22^. Using the Micro-Meta App, key microscope components and hardware specifications, including objectives, detectors, filters, light sources, acquisition modes, and calibration parameters, are systematically documented (Fig. 3). These details are essential for the accurate interpretation and reproducibility of microscopy data. Micro-Meta App supports tiered metadata reporting levels as defined by the 4DN-BINA-OME initiative ^36^, and exports metadata in JSON format, which is stored as an attachment to the corresponding experiment record in eLabFTW. Because microscope hardware specifications typically apply to the entire study, their metadata are annotated at the “Screen”, thereby avoiding redundancy across plates and wells. Practical guidance for implementing each metadata step is provided in the accompanying Jupyter Notebook on GitHub (Code Availability section).

### Data Handling

On the data-handling level within the workflow, **OMERO** serves as the central infrastructure for the storage, organization, and access of microscopy image data (Fig. 2). Raw image files from proprietary and open formats are ingested via Bio-Formats, along with their associated acquisition metadata. Images are uploaded through OMERO.insight and organized according to the experimental design using “Screens.” Screens are used for automation-assisted fluorescence imaging workflows and represent multi-well formats typical of automated microscopy. Within the workflow, “Screens” organize images into Plates and Wells, reflecting the physical laboratory setup for cell cultivation. This hierarchical structure (Screens → Plates → Wells → Images) provides a flexible, experiment-aligned data organization within OMERO, as shown in Fig. 4.

**Figure 4.**
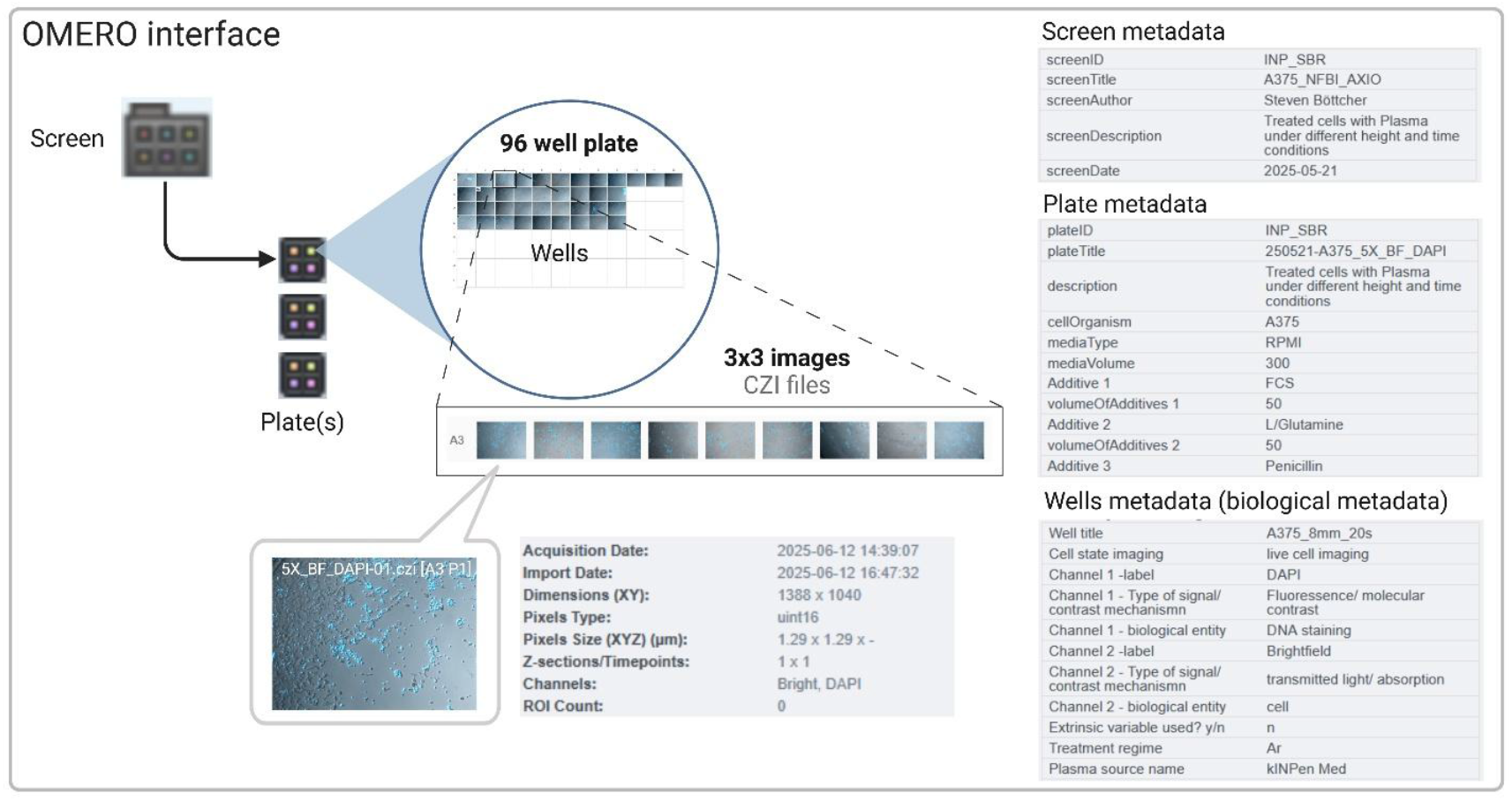
OMERO data organization and metadata annotation. (Left) Schematic representation of the OMERO data hierarchy used in automation-assisted fluorescence imaging. Experiments are organized into “Screens” comprising one or more multi-well Plates. Each “Plate” consists of individual “Wells”, and each well may contain multiple image fields (illustrated as a 3 × 3 tiled acquisition), reflecting automated microscopy workflows. (Right) Exemplary metadata annotations associated with the OMERO hierarchy, shown as key–value pairs (KVPs). Metadata can be assigned at different hierarchical levels (Screen → Plate → Well → Image) and are directly linked to the corresponding image data within the workflow. Created in BioRender. Bekeschus, S. (2026) https://BioRender.com/whuauio.

When images are uploaded via OMERO.insight, the associated acquisition metadata are automatically extracted and transferred to OMERO together with the images, named as “Original Metadata”. Within the workflow, experimental metadata at the “Screen”, “Plate”, and “Well” levels, previously created using Adamant and the Micro-Meta App and stored as JSON files in eLabFTW, are programmatically imported into OMERO. This integration is implemented using Python functions based on the omero-py and elabapi-python libraries, which are executed in the Jupyter notebook environment connecting OMERO and eLabFTW via their respective APIs. Users provide the “Screen”, “Plate”, and “Well” identifiers, as well as the corresponding eLabFTW entry IDs, through structured fields in the Jupyter notebook on GitHub (see Code Availability section). Access to eLabFTW records is controlled via user-specific API tokens.

One of the key advantages of this workflow step is the automated extraction and integration of metadata from different disciplinary contexts via their respective eLabFTW entries, followed by annotation at the appropriate organizational level within OMERO. As a result, metadata do not need to be manually exchanged between systems or annotated redundantly by multiple users. Instead, scientists working with image data gain access to consolidated metadata from biological experiments, plasma treatments, and microscopy acquisitions in a single, structured location aligned with OMERO’s organizational hierarchy. This process is illustrated in Fig. 5, which demonstrates how the Jupyter notebook interface enables users to annotate image data in OMERO by specifying screen names, plate names, and metadata identifiers retrieved from the eLabFTW. In this example, users enter metadata into predefined fields (Fig. 5) corresponding to OMERO’s organizational levels (“Screen”, “Plate”, and “Well”). Once populated, the notebook automatically establishes links between the eLabFTW metadata (e.g., experiment identifier, operator, and date) and the corresponding image containers in OMERO, ensuring consistent and reproducible metadata integration across the workflow.

**Figure 5.**
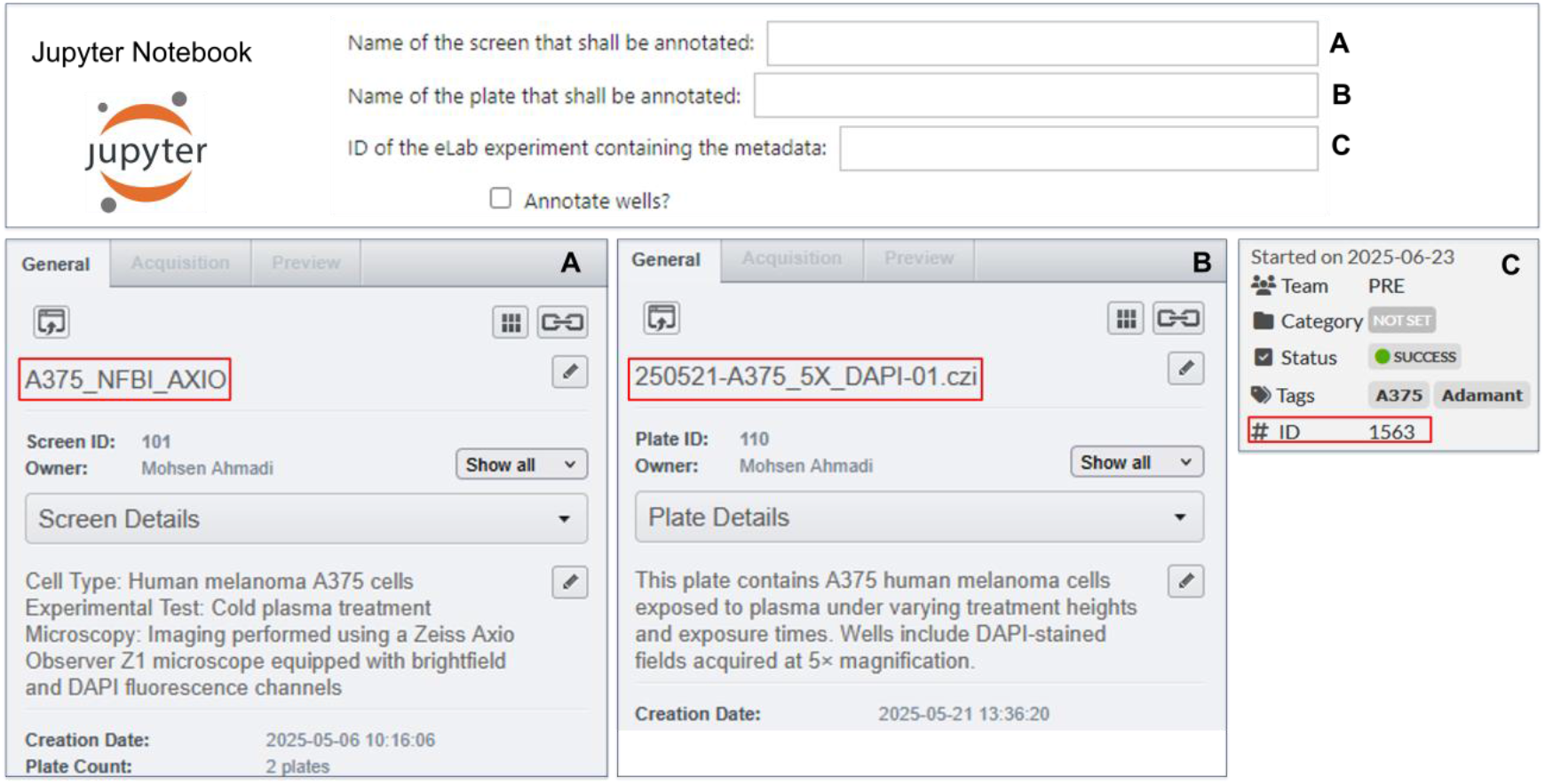
Jupyter notebook interface for image data annotation on OMERO at the “Screens”, “Plates”, and “Wells” levels. (A) input field for specifying the “Screen” name to be annotated in OMERO, (B) input field for specifying the “Plate” name within the selected screen, (C) field for entering the eLabFTW experiment ID, which links the OMERO dataset to the metadata recorded in eLabFTW.

After image annotation within the workflow, OMERO maps image acquisition metadata to the Open Microscopy Environment (OME) data model ^37^, translating file-specific parameters into a standardized structure. These parameters are stored as key-value pairs (KVPs) ^9^, which are required for the machine-readable annotation of each image and its experimental context. To incorporate domain-specific metadata generated in eLabFTW, Adamant, and the Micro-Meta App (see Metadata section), the workflow uses OMERO’s flexible annotation mechanisms, including tag-based and KVP annotations (Fig. 4). KVP annotations encode structured metadata as explicit attribute-value pairs, allowing biological, plasma-, and imaging-relevant parameters to be captured in a semantically consistent manner. For example, datasets are annotated with entries such as cell organism = A375, treatment regime = argon (Ar), cell treatment time [s] = 20, cell treatment distance [mm] = 8, and channel 1 - label = DAPI. In parallel, tag annotations are applied at the “Screen”, “Plate”, and “Well” levels to label datasets with key experimental attributes such as cell line, plasma treatment, or time point, enabling intuitive navigation and filtering across imaging studies.

### Data Processing

Image data and associated metadata are stored in OMERO (accessed by OMERO.web and OMERO.insight), enabling interactive use and programmatic downstream analysis. A key feature of the workflow is its interoperability with external analysis environments through graphical interfaces and APIs, supporting automated pipelines and exploratory analysis. Processed results, such as measurements generated with open-source image analysis tools ^38^, can be stored in OMERO as KVPs and/or an attached CSV file, establishing a closed loop between data acquisition, analysis, and reuse. Automated image analysis within the workflow combines technical expertise, scripting approaches (e.g., ImageJ/FIJI macros), and dedicated software components ^39^. In this context, OMERO integrates with image analysis tools, including CellProfiler for quantitative image analysis, FIJI/ImageJ for interactive and scripted processing, ilastik for machine-learning-based segmentation and classification, Orbit and QuPath for large-scale and whole-slide image analysis, and TrackMate for single-particle and cell tracking available at https://omero-guides.readthedocs.io/en/latest/external_tools.html.

Workflow-management environments such as KNIME ^20^, ImageJ integrated in KNIME ^40^, and Galaxy ^20^ can also be utilized to support automated and reproducible image analysis workflows. The choice of analysis tools depends on the specific experimental requirements and the user’s level of expertise. Python-based scripts can be used for custom image analysis and feature extraction, while FIJI is used for macro-based batch processing and interactive image analysis ^41^. All tools interact bidirectionally with OMERO-hosted data, enabling data retrieval and result storage (Fig. 2). Analysis results, including quantitative measurements, such as regions of interest (ROIs) and cell counts, can be stored as KVPs in OMERO for each image or linked back to experimental records in eLabFTW, for consistency between raw data, metadata, and analysis outputs.

In this section, we describe a FIJI-OMERO plugin recommendation in which the implemented infrastructure serves as the basis for downstream image analysis in FIJI via the corresponding plugin, illustrating how image data and metadata can be exchanged within the OMERO ecosystem to support interoperable analysis. The data processing level accommodates automated and interactive analysis strategies, tailored to experimental demands such as automation-assisted fluorescence imaging of multi-well plates or detailed single-image evaluation ^42^. Within this context, the FIJI-OMERO plugin (https://github.com/ome/omero-guide-fiji) is recommended for processing and managing annotated image datasets. This plugin establishes a secure connection to OMERO.insight server using user-specific authentication credentials that authorize access to image data and metadata (official OMERO plugin for ImageJ/FIJI is available at https://omero-guides.readthedocs.io/en/latest/fiji/docs/installation.html). Within FIJI, ROIs, such as areas containing dead cells or nuclei labeled with DAPI signals, can then be interactively defined or automatically detected and selected for quantitative analysis. In addition, ROIs can also be defined programmatically and analyzed using macros, which users write or customize to match their experimental needs. Once defined, a macro that referred to a scripted, automated sequence of image processing and analysis steps (e.g., ImageJ/FIJI) can be applied across datasets, retrieving images from OMERO and saving processed results, including images and data files such as CSV, back to the OMERO server. To support this, the OMERO plugin in FIJI provides functions including “Batch process”, “Connect to OMERO”, “Save image(s) or ROIs to OMERO”, “Save results to OMERO”, and “OMERO extensions” ^43^. FIJI/ImageJ macro workflows that access image IDs from OMERO and perform image processing, demonstrating practical batch analysis using OMERO macro extensions, are available via the German BioImaging GitHub repository: https://github.com/German-BioImaging/fiji_omero_workflows.

FIJI macros in the “OMERO Batch Plugin” implement processing pipelines for tasks in pre-processing and segmentation, including background subtraction, sharpening, particle analysis, and thresholding. Users process images stored on the OMERO server; the input section allows the selection of folders, loading of predefined ROIs, and recursive processing of subfolders. A task-specific FIJI macro file can be selected, which returns processed images, results tables, ROIs, or log files, all of which contribute to a transparent provenance trail.

### Example Use Case

To illustrate the practical application of the proposed workflow and to support its adoption by the community, this section demonstrates how the proposed workflow is applied in practice using a concrete **gas plasma-treated cell imaging dataset**. In this use case, adherent A375 melanoma cells are cultured in vitro and exposed to gas plasma (kINPen Med ^44^) under defined treatment conditions and subsequently subjected to automation-assisted ZEISS Axio Observer Z1. The example is structured according to the individual steps implemented by the Jupyter notebook on GitHub (Code Availability section), guiding the reader from data ingestion to metadata annotation and reuse. The dataset used for this demonstration is stored in the INPTDAT research data repository and serves as a documented reference implementation of the workflow (Data Availability section).

Fig. 6 illustrates the practical use of the presented workflow for plasma medicine bioimaging data management. The workflow begins by establishing an authenticated connection to OMERO and eLabFTW via their respective APIs. Within the Jupyter notebook, users provide their OMERO credentials and an eLabFTW API token, which are used to securely access image datasets and experimental metadata. This step is needed so that all subsequent operations (metadata retrieval, annotation, and validation) of the workflow are carried out within the given access permissions. Next, the user specifies the target OMERO objects to be annotated. In this example, a “Screen” representing an imaging experiment of cells is selected. The screen contains multiple “Plates” corresponding to 96-well imaging experiments, each comprising “Wells” that represent individual plasma-treated and untreated conditions. Metadata are captured and annotated at multiple OMERO hierarchical levels to reflect the experimental structure of a bioimaging study in plasma medicine with a persistent linkage between plasma treatment conditions and biological metadata (Table 2).

**Table 2.** Hierarchical “Screen”, “Plate”, and “Well” metadata for the example use case.

| Key | Value |
| --- | --- |
| **Screen (metadata collected with Admant)** |  |
| screenID | INP_SBR |
| screenTitle | A375_NFBI_AXIO |
| screenAuthor | Author(s) |
| screenDescription | Treated cells with plasma under different height and time conditions |
| screenDate | 2025-05-21 |
| **Plate (metadata collected with Admant)** |  |
| plateID | INP_SBR |
| plateTitle | 250521-A375_5X_DAPI |
| description | Treated cells with plasma under different height and time conditions |
| cellOrganism | A375 |
| mediaType | RPMI (Roswell Park Memorial Institute medium) |
| mediaVolume [μL] | 300 |
| Additive 1 | FCS (Fetal Calf Serum) |
| volumeOfAdditives 1 [μL] | 50 |
| Additive 2 | L/Glutamine |
| volumeOfAdditives 2 [μL] | 50 |
| Additive 3 | Penicillin |
| volumeOfAdditives 3 [μL] | 50 |
| Additive 4 | Streptomycin |
| volumeOfAdditives 4 [μL] | 50 |
| seedingDensity [cells/mm<sup>2</sup>] | 33.3 |
| layerDescription | Monolayer |
| plateDate | 2025-05-21 |
| creatorName | Author(s) |
| plateFeatures | Thermo Scientific NUNC Edge 96 well plate |
| imagingWithLid | True |
| microscopeName | Axio Observer Z1 |
| **Well (metadata collected with REMBI-compatible Excel template)** |  |
| Well title | A375_8mm_20s |
| Cell state imaging | Live cell imaging |
| Channel 1 - Label | DAPI |
| Channel 1 - Type of signal/contrast mechanisms | Fluorescence/ molecular contrast |
| Channel 1 - biological entity | DNA staining |
| Channel 2 - Label | Brightfield |
| Channel 2 - Type of signal/contrast mechanisms | Transmitted light/ absorption |
| Channel 2 - Biological entity | Cell |
| Extrinsic variable used? y/n | n |
| Treatment regime | Argon (Ar) |
| Plasma source name | kINPen Med |
| Cell treatment time [s] | 0, 20, 60, 180 |
| Cell treatment conditions | Room temperature and moisture |
| Cell treatment distance [mm] | 8, 12, 16, 20 |

**Figure 6.**
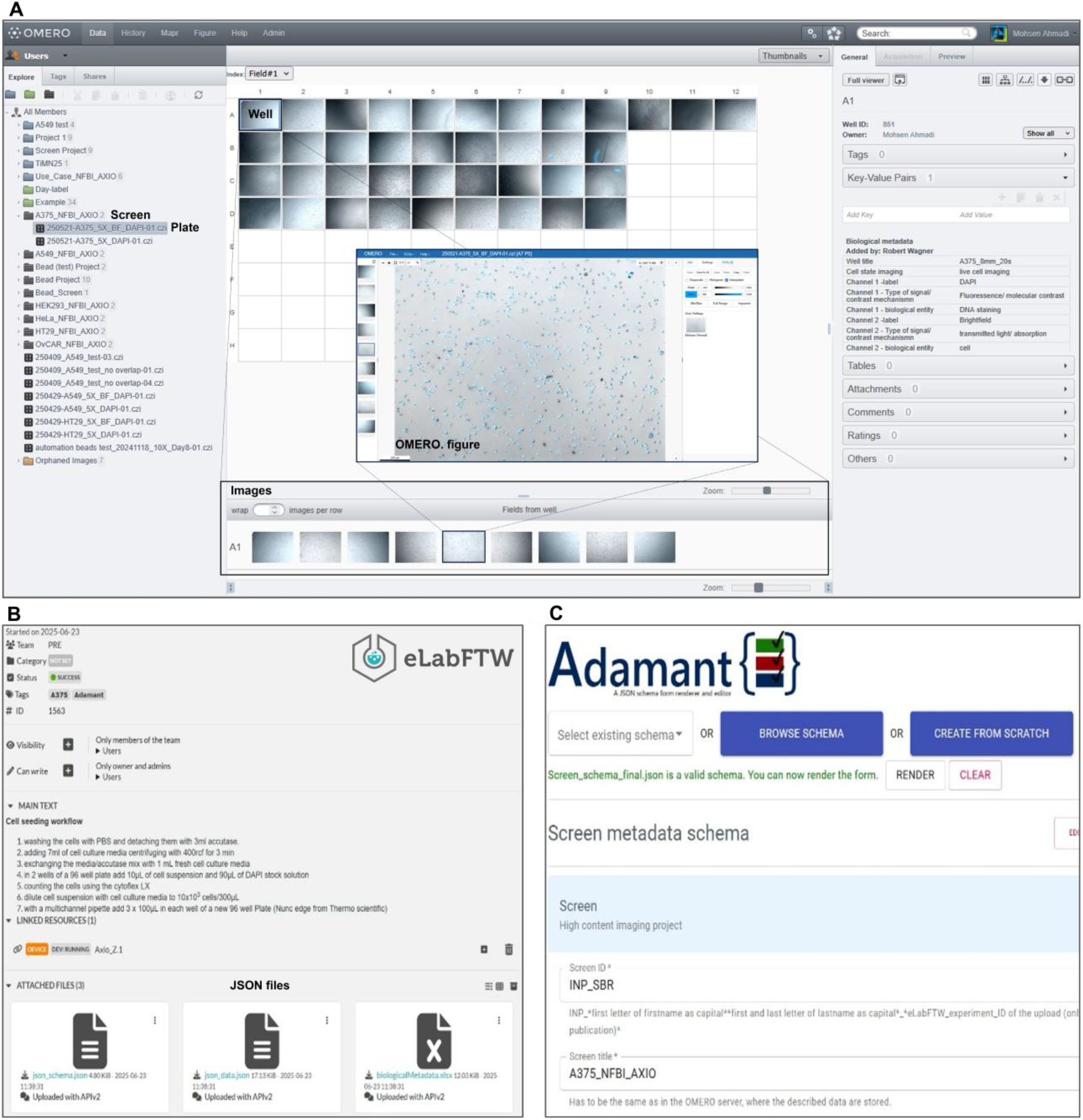
Example workflow in the OMERO user interface illustrating the dataset hierarchy, metadata panel, and preview window. (A) The OMERO interface displays an automation-assisted fluorescence imaging plate, including plate layout, well-level image previews, and a representative image field. Biological metadata associated with each well are shown in the metadata panel on the right, (B) An eLabFTW entry documenting the corresponding experimental procedure, linked resources, and attached JSON and CSV files that reference the imaging workflow, (C) The Adamant interface is used to define and validate the screen metadata schema. Full access to the dataset, including guiding documentation and all materials required to reproduce the example use case, is provided by INPTDAT at https://doi.org/10.34711/inptdat.1013.

These identifiers of the specific “Wells”, “Plates”, and “Screens” are entered once into predefined notebook fields and reused throughout the workflow (Fig. 6A). The notebook then retrieves structured experimental metadata from eLabFTW (Fig. 6B). These metadata are generated during the experiment using Adamant and include information on biological samples, plasma treatment parameters (e.g., plasma treatment time and distance), and experimental design (Fig. 6C). Metadata are stored in eLabFTW as JSON attachments linked to the corresponding experiment entry and are programmatically accessed via the ELN API. Retrieved metadata are validated against the expected schema and mapped to the OMERO organizational hierarchy. “Screen” metadata describes the overall study, “Plate” metadata captures experimental layouts, and “Well” metadata shows treatment-specific parameters (see Fig. 6A). In the annotation step, the notebook attaches selected metadata as KVPs (as represented in Table 2) to the OMERO hierarchy (“Screen”, “Plate”, and “Wells”) via provided scripts. These annotations link image data to their biological, plasma, and imaging context in a machine-readable form. Once completed, the image dataset is annotated, searchable, and reusable in OMERO, with plasma treatment parameters, biological context, and imaging metadata persistently linked to each image. The dataset can be accessed for downstream image analysis, enabling quantitative assessment and comparison of gas plasma-induced cellular responsesas as ROIs based on imaging data.

### FAIR Assessment

The RDM workflow for bioimaging in plasma medicine presented in this paper operationalizes the FAIR data principles across the entire bioimaging data lifecycle. Each FAIR dimension was systematically evaluated, as summarized in Table 1, to confirm that the workflow fulfils the corresponding requirements.

#### Metadata

The metadata level of the workflow fulfils the FAIR principles through the systematic use of discipline-appropriate, ontology-enriched metadata schemas and their consistent application across plasma treatments, biological assays, and microscopy imaging. Metadata are validated against JSON schemas and exported in a machine-readable JSON format, supporting interoperability through standardized representations (I1, Table 1) and ontology-based annotations aligned with community standards, such as REMBI for biological images ^29^ and ChEBI for chemical entities ^45^ (I2, R1, Table 1). Structured metadata from eLabFTW, Adamant, and the Micro-Meta App are consolidated into OMERO at the data handling level (see Fig. 2), where experimental records and image datasets are assigned unique identifiers to support findability and long-term accessibility (F1, A2, Table 1). The integration of biological metadata, plasma treatment parameters, and microscopy settings across systems establishes qualified references between metadata entities (I3, Table 1) and results in rich, contextualized datasets (F2, Table 1). Metadata and data stored in OMERO and exchanged in JSON format via eLabFTW-linked records ensure controlled accessibility (A1, Table 1) and traceability of the experimental context (F3, Table 1).

#### Data handling

Data annotation mechanism at the data handling level in OMERO provides the technical foundation for implementing FAIR principles across the workflow. OMERO assigns persistent internal identifiers to datasets ^9^ (e.g., “Screen” and “Plate” IDs; F1, Table 1), explicitly links metadata to the data they describe (F3, Table 1), and supports rich annotations for detailed dataset descriptions (F2, Table 1). All datasets and associated metadata are indexed and searchable using the OMERO interface (F4, Table 1). Furthermore, the workflow uses OMERO’s standardized access mechanisms, including its open API and web interface, to enable metadata and data retrieval (A1, Table 1). Access is provided in a controlled manner, typically by request and permission, while remaining widely implementable across systems (A1.1). User authentication and project-based access control are applied where required to protect sensitive data (A1.2). Importantly, within the workflow, metadata remain accessible in OMERO even if the associated image data are archived or removed, thereby supporting long-term metadata availability (A2, Table 1). In terms of interoperability, the workflow uses OMERO support for machine-readable metadata formats such as JSON and XML (I1, see Table 1). In addition, the workflow establishes qualified references between related metadata entities within OMERO, supporting semantic linking across experimental components (I3, Table 1). Finally, the workflow reusability is ensured by the rich description of experimental and technical metadata in OMERO (R1, Table 1), the ability to annotate datasets with licenses or data use terms (R1.1), provenance tracking via user logs and timestamps (R1.2), and alignment with relevant community standards (R1.3).

#### Data processing

At the data processing level, the FIJI-OMERO integration operationalizes the FAIR principles. Secure and user-specific authentication provides authorized access to image data and metadata stored in OMERO (A1, A1.2, Table 1). ROIs can be defined interactively or programmatically, and analysis steps are implemented as reusable and customizable macros aligned with experimental needs (R1, R1.2, Table 1). These macros support scalable batch processing of OMERO-hosted datasets, retrieving images via APIs, and returning results in a structured and machine-readable format (A2, I1, Table 1). Processed outputs, including measurements, ROIs, and logs, are written back to OMERO as annotations, preserving links to the original data and enabling traceable provenance (I2, R1.3, Table 1). This establishes a closed, reproducible processing loop that supports standardized analysis, transparent documentation, and FAIR-compliant data reuse.

#### Overall FAIR assessment of the workflow

Taken as a whole, the workflow demonstrates a coherent and end-to-end implementation of the FAIR principles across metadata capture, data handling, and data processing. Findability is set through persistent internal identifiers, structured organizational hierarchies, and comprehensive indexing that allow metadata and image data to be reliably located and referenced throughout their lifecycle. Accessibility is supported by standardized network-based access mechanisms combined with authentication and authorization controls, while preserving metadata availability beyond the lifetime of the raw image data. Interoperability is achieved at the technical level through the consistent use of machine-readable formats, standardized schemas, and explicit semantic links between related entities, providing a stable foundation for cross-system exchange and integration. Reusability is enabled by the rich contextualization of data using community-aligned metadata standards and by maintaining traceable links between experiments, images, and derived results. In the FAIR assessment of the workflow, the Jupyter Notebook located on GitHub contains all required tools for data curation and annotation and serves as an executable documentation layer. It makes the workflow steps reproducible, links data and metadata across OMERO, eLabFTW, and Adamant via APIs, implements standardized schemas in machine-readable form, and records provenance, parameters, and versions. While not a repository itself, it implements **FAIR compliance** by operationalizing, validating, and demonstrating the workflow in a reusable way.

While certain aspects, such as vocabulary refinement and extended provenance tracking, remain subject to iterative improvement, the workflow as implemented already constitutes a FAIR-compliant and extensible framework for bioimaging data management in plasma medicine. FAIR principles define high-level objectives for data management but do not provide detailed, domain-specific guidance on how these objectives should be implemented in practice. Therefore, the **DSW FAIR assessment** was used to determine which components of the workflow contribute to overall FAIR compliance ^46^. Using the Common DSW Knowledge Model (see Method section), the structured assessment illustrates the workflow’s FAIR compliance levels and the contribution of specific questionnaire topics to each FAIR section (Fig. 7).

**Figure 7.**
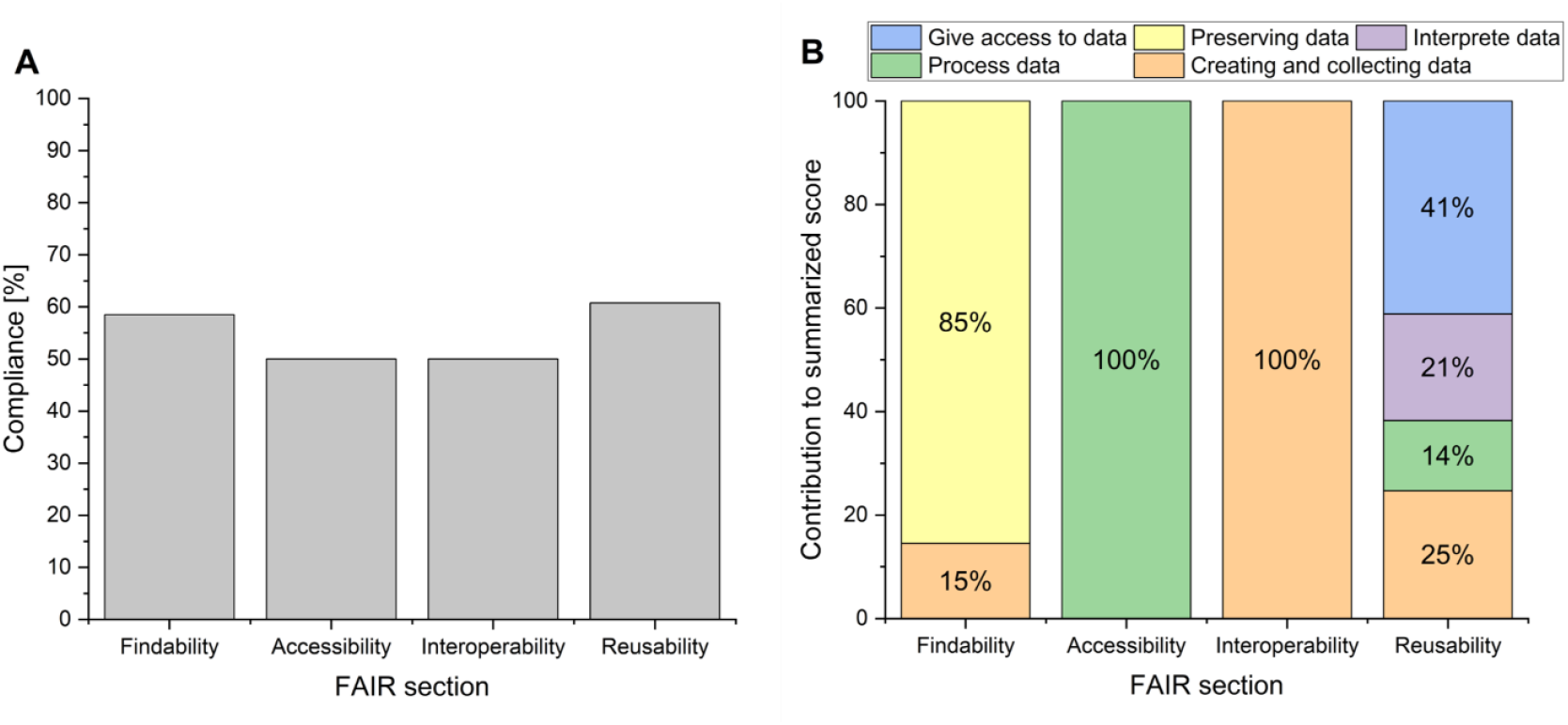
Summary report generated with the Data Stewardship Wizard using the Common DSW Knowledge Model 2.6.10. (A) average FAIR compliance scores of the implemented data management workflow for bioimaging in plasma medicine; (B) contribution of selected questionnaire domains to the summarized FAIR scores. The corresponding knowledge model is available at: https://registry.ds-wizard.org/knowledge-models/dsw:root:2.6.12.

Fig. 7A shows the compliance scores for each FAIR section, indicating moderate fulfilment of findability, accessibility, interoperability, and reusability. The limitations in the area of accessibility are primarily because access to stored microscopy images, i.e., particularly those involving patient samples from partner hospitals, must be individually regulated. As a result, full accessibility compliance cannot be fully achieved until user-specific access requirements are more clearly defined. Fig. 7B breaks down how different activity domains within the workflow contribute to these scores. The findability score is primarily driven by data creation and collection, whereas accessibility and Interoperability are fully supported by processes related to giving access to data and preserving data. Reusability, in contrast, is influenced by a broader combination of activities, including data interpretation, data creation and collection, data processing, and data preservation, reflecting its dependence on complete and semantically rich metadata. This variation originates from the questionnaire’s adaptive structure; depending on prior responses, different follow-up questions may be triggered or omitted, leading to workflow-specific differences in assessment depth and emphasis.

### Conclusion and Outlook

We introduced a data management workflow for bioimaging in plasma medicine that harmonizes image acquisition, experimental (gas plasma and biological experiments) documentation, and metadata annotation of the bioimaging data lifecycle across OMERO, eLabFTW, Adamant, and the Micro-Meta App, with all components interconnected via APIs within a Jupyter notebook environment. Within OMERO, the workflow results in well-structured image datasets enriched with linked experimental and technical metadata, including plasma treatment parameters, biological assay information, and microscopy settings. Images are organized in “Screens”, “Plates”, and “Wells” and annotated with standardized metadata and identifiers as key-value pairs (KVPs). By systematically linking imaging data to experimental and technical metadata, the workflow provides end-to-end traceability from experiment data standardization to reproducibility and reuse. Interoperability is achieved through established community standards for image formats (e.g., CZI, ND2, LIF, OME-TIFF, and OME-NGFF) ^47^ and JSON-based metadata exchange, with APIs and controlled import-export mechanisms preserving metadata integrity. Furthermore, semantic interoperability is strengthened through controlled vocabularies and community models, notably the OME data model ^37^ and the Plasma-MDS ^31^ extension. The resulting machine-actionable outputs permit downstream applications, including automated image analysis and AI-driven methods for automation-assisted fluorescence imaging of multi-well plate experiments in plasma medicine. Despite strong FAIR alignment, some challenges remain. Metadata harmonization across imaging platforms still requires manual curation to align heterogeneous formats, terminology, and parameter definitions across different acquisition systems. On the other hand, consistent long-term archiving with public repositories (e.g., IDR and BIA) is under active development within the OMERO ecosystem. Ongoing efforts in initiatives such as NFDI4Bioimage and Plasma-MDS are expected to address these limitations, further extending the applicability and sustainability of the workflow not only in plasma medicine but also in bioimaging and biomedical research domains.

## Data availability

The dataset including additional guiding documentation and all materials required to reproduce the example use case, is available via INPTDAT (https://doi.org/10.34711/inptdat.1013). The included image database contains all raw images in.czi and TIFF/XML format.

## Code availability

The Jupyter notebook-based implementation of the workflow is publicly accessible via the project’s GitHub repository (https://github.com/INP-SDT/Bioimaging-Data-Management-Workflow-for-Plasma-Medicine).

## Acknowledgements

This work is funded by the Deutsche Forschungsgemeinschaft (DFG, German Research Foundation) under the National Research Data Infrastructure – NFDI 46/1 – 501864659. The authors acknowledge Steven Böttcher for his efforts in conducting the biological assays and managing the sample treatments. Several graphics within Figures 1 to 4 were created using a commercial license of BioRender.

## Author contributions

M.A. wrote the manuscript, designed and implemented the workflow, and performed data collection and integration. R.W. revised and edited the manuscript and contributed to workflow implementation, Jupyter Notebook development, and code writing. S.B. contributed expertise in plasma medicine and supported the data design and manuscript revision. M.M.B. contributed to the conceptual design, data interpretation, and manuscript revision.

## Competing interests

The authors declare no competing interests.

## Notes

https://doi.org/10.34711/inptdat.1013

https://github.com/INP-SDT/Bioimaging-Data-Management-Workflow-for-Plasma-Medicine

### Competing Interest Statement

The authors have declared no competing interest.

